# Utilization of Arabidopsis E3 ubiquitin decoys high-throughput yeast screen platform to dissect the ubiquitin-mediated circadian clock regulation

**DOI:** 10.1101/2025.08.31.673341

**Authors:** Yi-Tsung Tu, Chen-An Chen, Joshua M. Gendron, Chin-Mei Lee

**Affiliations:** Institute of Plant Biology, National Taiwan University, Taipei, 106319, Taiwan; Molecular, Cellular and Developmental Biology, Yale University, New Haven, CT, 06511, USA; Department of Life Science, National Taiwan University, Taipei, 106319, Taiwan; The Master Program in Global Agriculture Technology and Genomic Science, National Taiwan University, Taipei, 106319, Taiwan; International Joint Degree Master’s Program in Agro-biomedicine in Food and Health, National Taiwan University, Taipei, 100233, Taiwan

**Keywords:** E3 ubiquitin ligase, F-box protein, U-box protein, ubiquitination, circadian clock, PUB18, JMJD5, LHY

## Abstract

Protein ubiquitination, mediated by E3 ubiquitin ligases, is a critical regulatory mechanism of eukaryotic cellular processes, including circadian clock function. However, identifying E3–substrate pairs remains technically challenging due to substrate instability and the genetic redundancy of E3s. To overcome these limitations, we developed a high-throughput yeast two-hybrid E3 decoy screening platform, enabling systematic mapping of E3–substrate interactions. Using a library of 283 Arabidopsis F-box and U-box E3 decoys, we screened 21 core circadian clock regulators and identified 77 potential E3–substrate interaction pairs involving 56 E3s and 16 clock proteins. Focusing on high-confidence hits, we demonstrated that PUB18 physically interacts with the central clock regulators LHY and JMJD5 and promotes their ubiquitination in planta. Genetic analyses further revealed that PUB18 and its homolog PUB19 function redundantly in circadian clock regulation. This study establishes the E3 decoy yeast two-hybrid platform as a versatile and scalable tool for dissecting ubiquitination networks in broad biological processes.

## Introduction

Protein ubiquitination is a critical post-translational modification that regulates diverse cellular processes in eukaryotes, including proteasome-mediated protein degradation, protein localization, transcriptional regulation, and signal transduction (Buetow and Huang, 2016). This process involves a cascade of enzymatic activities requiring three key enzymes: E1 ubiquitin-activating enzymes, E2 ubiquitin-conjugating enzymes, and E3 ubiquitin ligases (Vierstra, 2009). Ubiquitin is first activated by E1, transferred to E2, and ultimately conjugated to target proteins through the action of E3 ligases, which determine substrate specificity by bridging E2 enzymes with specific targets.

In Arabidopsis, the genome encodes two E1s, 37 E2s, and approximately 1400 E3 ligases, highlighting the extensive regulatory potential of E3s in plant biological processes (Stone, 2014). E3 ligases are classified into single-subunit types—such as HECT (homologous to E6-AP carboxyl terminus), RING (Really Interesting New Gene), and U-box—and multi-subunit complexes, including cullin-RING ligases (CRLs) such as SCF (SKP1–CUL1–F-box), BTB (Bric-a-brac-Tramtrack-Broad complex), DDB (DNA Damage-Binding), and APC (anaphase-promoting complex) (Vierstra, 2009). The large number of E3s and some of them modulating the same substrate often results in genetic redundancy. Additionally, many E3s promote degradation of their substrates, isolating E3–substrate interactions biochemically and genetically remains technically challenging.

To overcome these obstacles, the “E3 decoy” strategy was developed (Lee and Feke et al., 2018). In this approach, E3 constructs lacking the F-box or U-box domain cannot recruit E2 enzymes, preventing substrate ubiquitination while preserving substrate binding. As a result, substrates are stabilized, facilitating interaction studies. Feke et al. (2019) created a library of 176 F-box and 45 U-box E3 decoys in Arabidopsis to screen for phenotypes related to circadian rhythms and flowering time. While several E3 ligases were identified as candidate regulators in these processes (Feke et al., 2019; 2020), the platform had limitations. First, the space and labor demands restricted the environmental conditions that could be tested during phenotypic screening. Second, phenotypic variability could obscure subtle E3-dependent traits. Finally, identifying the direct substrates of E3 ligases remained challenging and required complementary techniques such as yeast two-hybrid (Y2H) assays or immunoprecipitation coupled with mass spectrometry.

To overcome these limitations and systematically identify E3 ligases involved in circadian clock regulation, we developed a high-throughput yeast two-hybrid E3 decoy screening platform. The circadian clock is an endogenous timekeeping system that synchronizes physiological and developmental processes with daily environmental changes, thereby enhancing organismal fitness (Dodd et al., 2005; Greenham and McClung, 2015). It operates through interlocked transcriptional and translational feedback loops and is entrained by environmental signals such as light and temperature (Nohales and Kay, 2016). In Arabidopsis, the MYB transcription factors CIRCADIAN CLOCK ASSOCIATED 1 (CCA1) and LATE ELONGATED HYPOCOTYL (LHY) peak at dawn and bind to Evening Element (EE) motifs to repress *TIMING OF CAB2 EXPRESSION1* (*TOC1*). TOC1, in turn, represses *CCA1* and *LHY* in the evening. TOC1 also represses morning-phased *PSEUDO-RESPONSE REGULATORS* (*PRR9, PRR7*, and *PRR5*) and the evening-expressed LUX ARRHYTHMO (LUX), which functions with EARLY FLOWERING 3 (ELF3) and ELF4 as the Evening Complex (Gendron et al., 2012; Huang et al., 2012). Beyond these feedback transcription-translation regulations, the histone demethylase JUMONJI DOMAIN-CONTAINING 5 (JMJD5) also represses *CCA1* and *LHY* and is reciprocally regulated by them (Jones et al., 2010; Lu et al., 2011).

Post-translational regulation by ubiquitination is critical for robust circadian function. The F-box protein ZEITLUPE (ZTL) acts as an evening-phase E3 ligase, targeting TOC1 and PRR5 for degradation in darkness (Mas et al., 2003; Kiba et al., 2007), with its maturation dependent on HEAT SHOCK PROTEIN 90 (HSP90) and GIGANTEA (GI) (Kim et al., 2011; Cha et al., 2017). Another E3 ligase, CONSTITUTIVE PHOTOMORPHOGENESIS 1 (COP1), targets ELF3 and GI for proteasomal degradation, with ELF3 acting as a substrate adaptor (Yu et al., 2008). Despite these insights, only a limited number of E3 ligases involved in circadian regulation have been characterized to date.

In this study, we constructed a yeast two-hybrid E3 decoy library comprising 283 F-box and U-box E3 ligases—covering approximately 37% of those encoded in Arabidopsis—and used it to systematically screen for interactions with 21 known circadian clock regulators. We identified 77 E3– clock regulator interaction pairs. Among these, we validated that PUB18 interacts with and ubiquitinates both LHY and JMJD5 in planta. Furthermore, we showed that PUB18 and its homolog PUB19 function redundantly in clock regulation. This high-throughput E3 decoy screening platform overcomes technical bottlenecks in identifying E3–substrate interactions and provides a versatile tool for studying ubiquitin-mediated regulation in the circadian clock and beyond.

## Results

### Construction of the E3 decoy library for a high-throughput yeast screen platform

To systematically identify potential E3 ubiquitin ligases (E3) that interact with clock regulators, we constructed a high-throughput yeast screen platform from the previously established E3 decoy collections by Feke et al. (2019). This E3 decoys collection contains 238 F-box type and 45 U-box type E3s with deletion of conserved F-box or U-box domains in the Gateway *pENTR* vectors, which facilitate the Gateway yeast expression vector pDEST32 by recombination. The deletions of the F-box or U-box domains in these E3 decoys eliminate the ubiquitination of their interacting substrates and thus stabilize them, which may reduce the false negative rates during the yeast screen. We successfully subcloned all F-box and U-box E3 decoys into the pDEST32 to generate GAL4 DNA-binding domain fusion E3 decoys library (or called pDEST32-E3 decoy library thereafter) and transformed them individually into the Y187 yeast strain (**Figure 1, Supplemental Table 1**). We grew and stored each clone in three 96-well plates to facilitate the high-throughput screen process.

**Fig 1.**
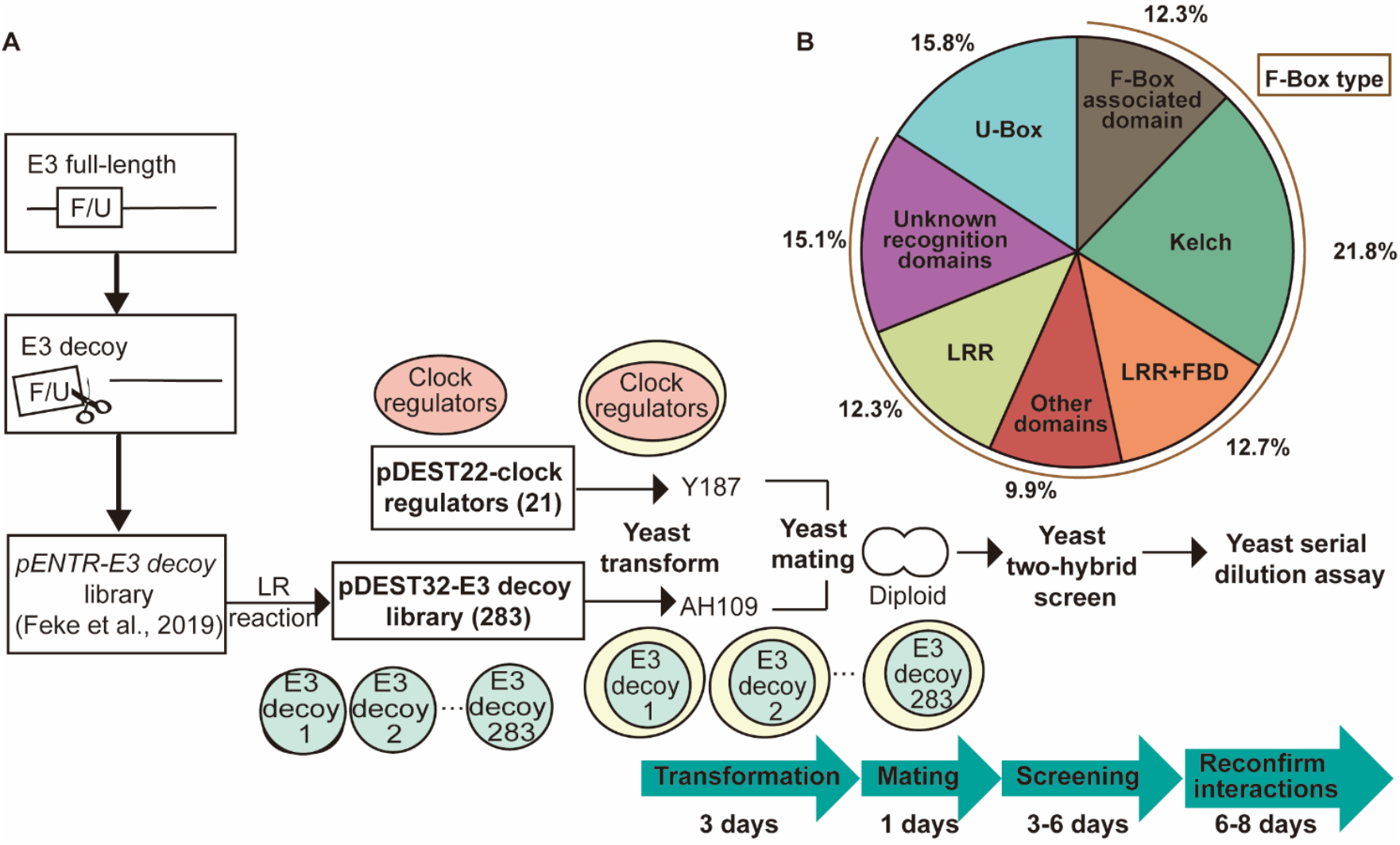
The construction of Arabidopsis E3 decoy yeast library and the process of high-throughput screening with circadian clock regulators. (**A**) The *pDEST32-E3 decoy* library was subcloned from 283 *pENTR-E3 decoy*s (Feke et al., 2019) and transformed into yeast AH109 strain individually to generate three 96-well plates as a high-throughput yeast screen platform. The 21 selected clock regulators (Supplemental Table 2) were cloned to *pDEST22* and transformed into Y187 strain. The diploid cells expressing E3 decoys and clock regulators were generated by yeast mating in the 96-well plates and selected on the SD-Leu/Trp and SD-Leu/Trp/His plates for interactions. This process takes about 7∼10 days. The positive interactions were selected based on the growth of colonies compared to the combination of the *pDEST22-clock regulator* and *pDEST32* empty vector. The diploid cells were further examined through serial dilution assays to confirm the selected positive interacting pairs, which take about 6 to 8 days. (**B**) The *pDEST32-E3 decoy* library consists of 45 U-box E3 decoys and 238 F-box E3 decoys (Supplemental Table 1). The 238 F-box E3 decoys include subfamilies with different conserved protein domains, such as F-box domain (FBD), Leucine-Rich Repeat (LRR), and the kelch domain.

The pDEST32-E3 decoy yeast library contains 45 U-box decoys, representing 70% of Arabidopsis’s total U-box E3 ligase (Yee and Goring, 2009). The U-box E3 decoys comprise 15.8% of the yeast library. The rest of them are F-box type E3 decoys, including F-box associated domain (12.3%), Kelch (21.8%), LRR and FBD (12.7%), LRR (12.3%), other domains (9.9%), and unknown recognition domain (15.1%) subfamilies (**Figure 1, Supplemental Table 1**) These 239 F-box E3 decoys represent 34% of the Arabidopsis F-box protein family (Gange et al., 2002).

### Screening for the E3s interacting with the clock regulators

We applied this E3 decoys yeast screen platform to identify the E3s interacting with 21 known clock regulators, including CCA1, LHY, PRR3, PRR7, PRR9, TOC1, REVEILLE 4 (RVE4), RVE6, RVE8, BROTHER OF LUX ARRHYTHMO (BOA/NOX), JMJD5, FLOWERING BHLH 1 (FBH1), TEOSINTE BRANCHED 1-CYCLOIDEA-PCF20 (TCP20), TCP22, HEAT SHOCK TRANSCRIPTION FACTOR B2b (HSFB2b), GI, LIGHT-REGULATED 1 (LWD1), LWD2, CCA1 HIKING EXPEDITION (CHE), NIGHT LIGHT–INDUCIBLE AND CLOCK-REGULATED 1 (LNK1), and CK2 (**Supplemental Table 2**). Most of these clock regulators are expressed rhythmically and peak at different times of the day (Wang and Tobin, 1998; Matsushika et al., 2000; Rawat et al., 2011; Dai et al., 2011; Jones et al., 2010; Nagel et al., 2014; Giraud et al., 2010; Pruneda-Paz et al., 2009; Kolmos et al., 2014; Park et al., 1999; Wu et al., 2016; Xie et al., 2014; Sugano et al., 1998). Moreover, they are responsible for regulating the circadian clock under different environmental signals, such as light and temperature (Nagel et al., 2014; Xie et al., 2014; Wu et al., 2016). These characteristics make condition-specific phenotypic analysis or biochemical identification a challenge for the E3-clock regulator pairs.

To set up the high-throughput yeast screen, we subcloned these 21 clock regulators into the *pDEST22* vectors to produce GAL4 activation domain fusion proteins and then transformed them into the yeast strain Y187, individually (**Figure 1**). We performed yeast mating to generate diploid cells by adding the Y187 yeast expressing *pDEST22*-clock regulator to the 96-well plates, in which each well contained AH109 yeast cells expressing one *pDEST32*-E3 decoy. The diploid yeast cultures were dropped onto the synthetic dropout (SD) medium without Leucine and Tryptophan (SD-Leu-Trp) for testing the expression of bait and prey vectors, and the medium without Leucine, Tryptophan, and Histidine (SD-Leu-Trp-His) for determining interactions. Our yeast screen platform and pipeline provide a high-throughput and cost-effective strategy to systematically identify the potential E3s regulating the circadian clock within 2 weeks to surpass genetic and biochemical challenges.

With this strategy, we examined 5943 interaction pairs and finally identified 77 potential interactions among the 56 E3 decoys and 16 clock regulators (**Figure 2**; **Supplemental Table 3**). In our screens, we did not find E3 interactors with PRR7, TCP20, LWD2, TCP22, and CHE. The previously reported interactions, such as GI-ZTL (Cha et al., 2017), were found. Additionally, two E3 decoys, COP9 INTERACTING F-BOX KELCH 1 COP9 INTERACTING F-BOX KELCH 1 (CFK1) and ALTERED CLOCK F-BOX 1 (ACF1), altered the phase of *CCA1* under circadian conditions (Feke et al., 2019), were found to interact with PRR9 and RVE6 in our screen, respectively (**Figure 2**; **Supplemental Table 3**).

**Figure 2.**
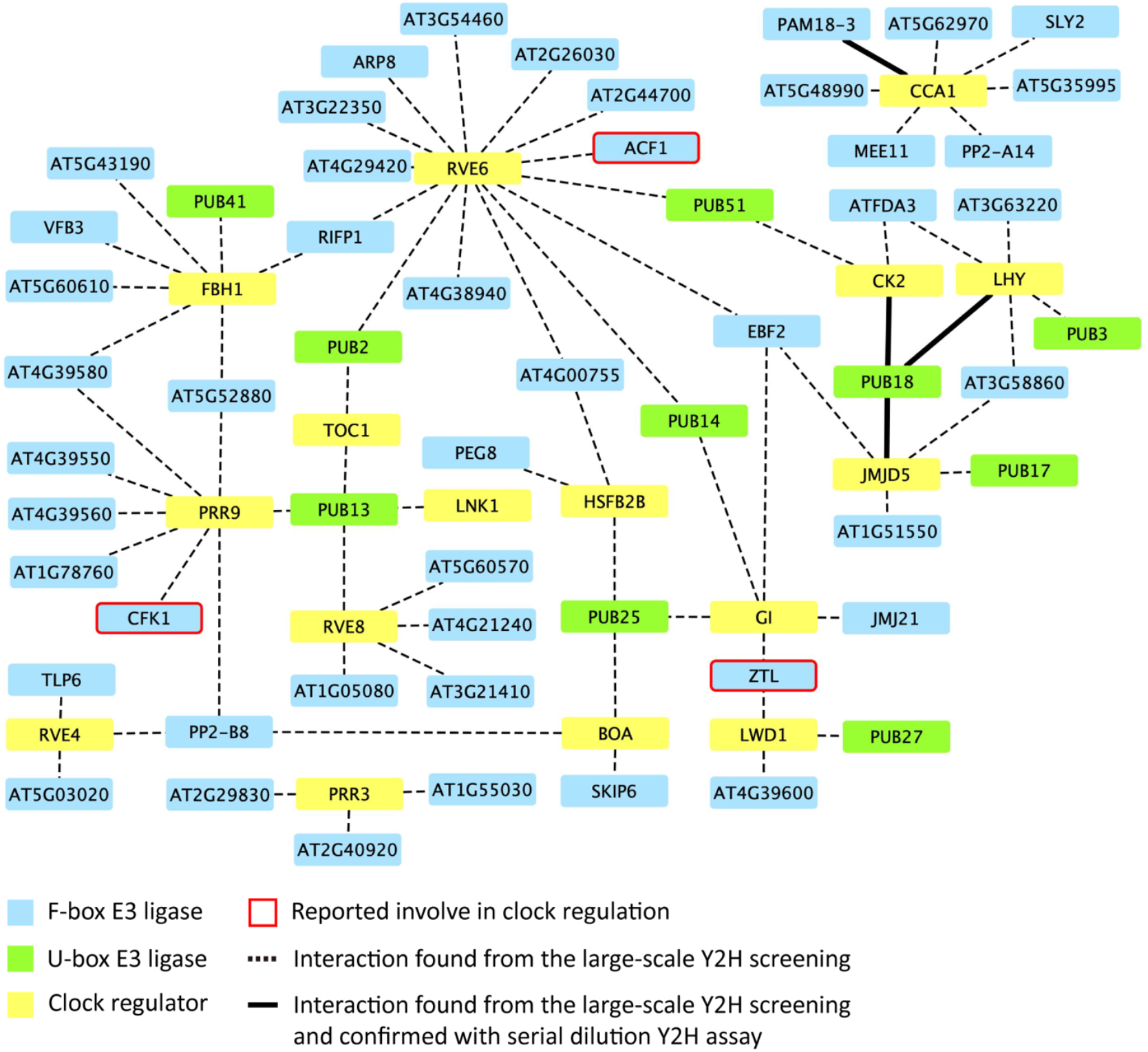
The E3 decoy-clock regulator interacting network from the high-throughput yeast hybrid screen. The protein-interacting network constructed from the high-throughput yeast hybrid screen illustrated the clock regulators (yellow box) and their interacting U-box (green box) and F-box (blue box) type E3 decoys with a dotted line. The interactions between E3 decoy candidates and CCA1, LHY, JMJD5, and CK2 were confirmed with yeast serial dilution assays and indicated by the solid lines. The E3 ubiquitin ligases previously reported to be involved in clock regulation, such as ZTL, ACF1, and CFK1, were labeled with red rims (Kim et al., 2007; Feke et al., 2019).

Among these potential interacting pairs, we chose to further verify the interactions from the high-throughput screen. We centered on the morning-expressed CCA1 and LHY (the right upper corner of **Figure 2**). Additionally, we included CK2 and JMJD5 as they share some E3-interacting candidates with CCA1 and LHY. We performed serial dilution assays to confirm the 20 interactions among CCA1, LHY, CK2, and JMJD5 with the potential E3 decoys from the initial high-throughput screen (**Figure 3; Supplemental Table 3**). The PRESEQUENCE TRANSLOCASE-ASSOCIATED MOTOR 18-3 (PAM18-3) decoy can interact with CCA1 (**Figure 3A**, top panel). Furthermore, the PLANT U-BOX 18 (PUB18) decoy could interact with LHY, JMJD5, and CK2 in yeast (**Figure 3B**). These results indicated about 20% hit rate in our high-throughput screening for the four clock regulators. This pilot experiment showed the effectiveness of narrowing down the interacting candidates within 6-8 days.

**Fig 3.**
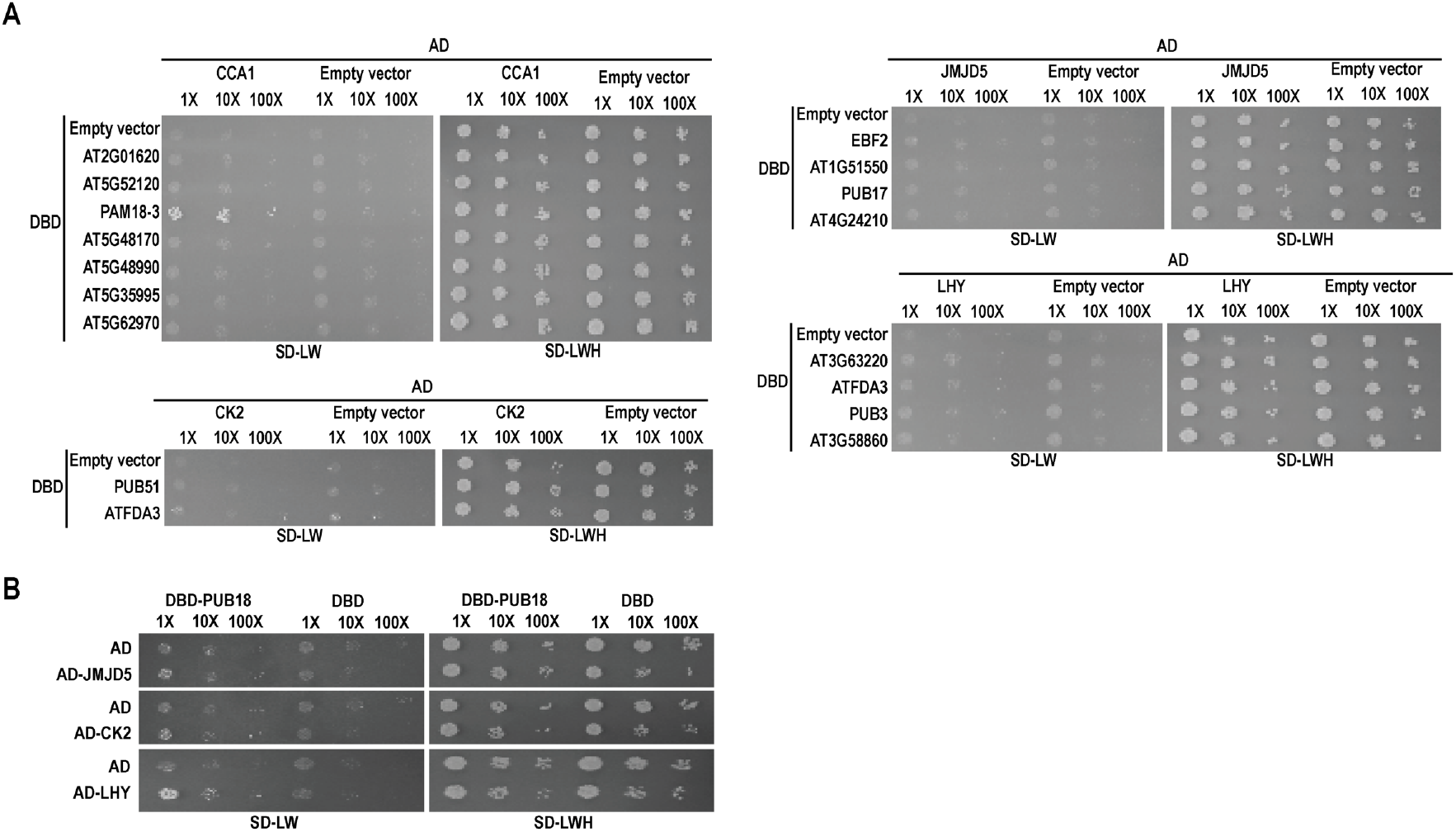
The results of yeast serial dilution assays confirmed the interactions between CCA1, LHY, JMJD5, and CK2 with E3 ubiquitin ligases. (**A**) The yeast two-hybrid assays of CCA1, LHY, JMJD5, and CK2 with their potential E3 decoy candidates from the high-throughput screen (**Figure 2**). The 10-fold (10X) and 100-fold (100X) dilutions were made from the yeast culture with O.D. 0.4 (labeled with 1X). The yeast culture was grown on SD-Leu-Trp (SD-LW) or SD-Leu-Trp-His (SD-LWH) medium for interaction tests. (**B**) The yeast hybrid assays confirmed that PUB18 interacts with LHY, JMJD5, and CK2.

### PUB18 interacts and ubiquitinates JMJD5 and LHY in plant

We next verified the interactions between PUB18 decoys or called PUB18dU, which lacks of U-box domain, with JMJD5, LHY, and CK2 in plant cells by performing bimolecular fluorescence complementation (BiFC) assay in *Nicotiana bethamiana*. The LHY interacted with PUB18dU in the nucleus, and the interactions of JMJD5-PUB18dU were detected in both the cytoplasm and the nuclei (**Figure 4A, left panel**). Previous studies showed that PUB18 localizes in the cytoplasm (Seo, et al., 2016), and JMJD5 localizes in the nucleus and cytoplasm (Lu et al., 2011). Therefore, PUB18dU and JMJD5 can interact in the nucleus and cytoplasm. However, PUB18dU did not interact with CK2 in plant. We further examined the interactions with full-length PUB18 (PUB18FL) and got similar results of PUB18dU (**Figure 4A, right panel**). To further corroborate the interactions of LHY-PUB18 and JMJD5-PUB18, we performed coimmunoprecipitation (Co-IP) assay. Both PUB18FL and PUB18dU can be pulled down by JMJD5 or LHY (**Figure 4B**). Taken together, these experiments supported that LHY and JMJD5 interact with PUB18 in plant cells.

**Figure 4.**
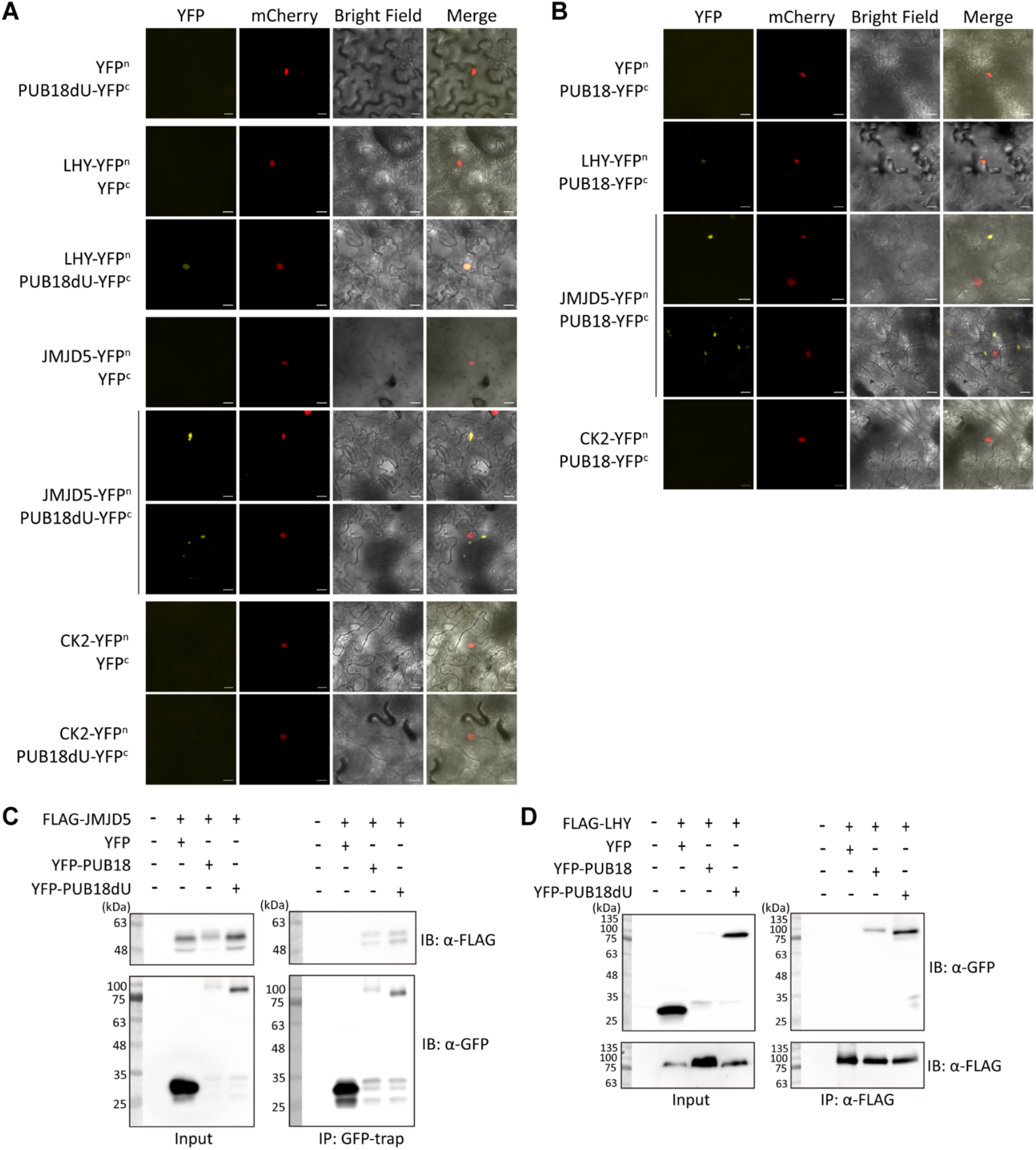
JMJD5 and LHY interact with PUB18 and PUB18dU in *N. benthamiana*. The Bimolecular fluorescence complementation (BiFC) assays to examine the interactions between JMJD5 and LHY fused to the N-terminal of YFP (YFPn) and **(A)** U-box-deleted PUB18 (PUB18dU) or **(B)** full-length PUB18 (PUB18) fused to the C-terminal of YFP (YFPc) in tobacco leaves. The NLS-mCherry was co-expressed as a nuclear marker. Scale bar= 20µm. The co-immunoprecipitation (co-IP) assays are used to determine the interactions between PUB18 and **(C)** JMJD5 or **(D)** LHY. The FLAG-JMJD5 or FLAG-LHY was co-expressed with PUB18 or PUB18dU fused to YFP tobacco leaves. GFP-Trap and anti-FLAG were used to immunoprecipitate YFP-PUB18/PUB18dU and FLAG-LHY, respectively. Anti-GFP and anti-FLAG antibodies were used for immunoblot analysis.

PUB18 was reported as a functional E3 (Seo, et al., 2016); we, therefore, determined whether PUB18 can ubiquitinate JMJD5 and LHY. Coexpression of PUB18 with either JMJD5 or LHY resulted in increased ubiquitination of JMJD5 and LHY (**Figure 5**). These results demonstrated that PUB18 is the E3 of JMJD5 and LHY clock regulators.

**Figure 5.**
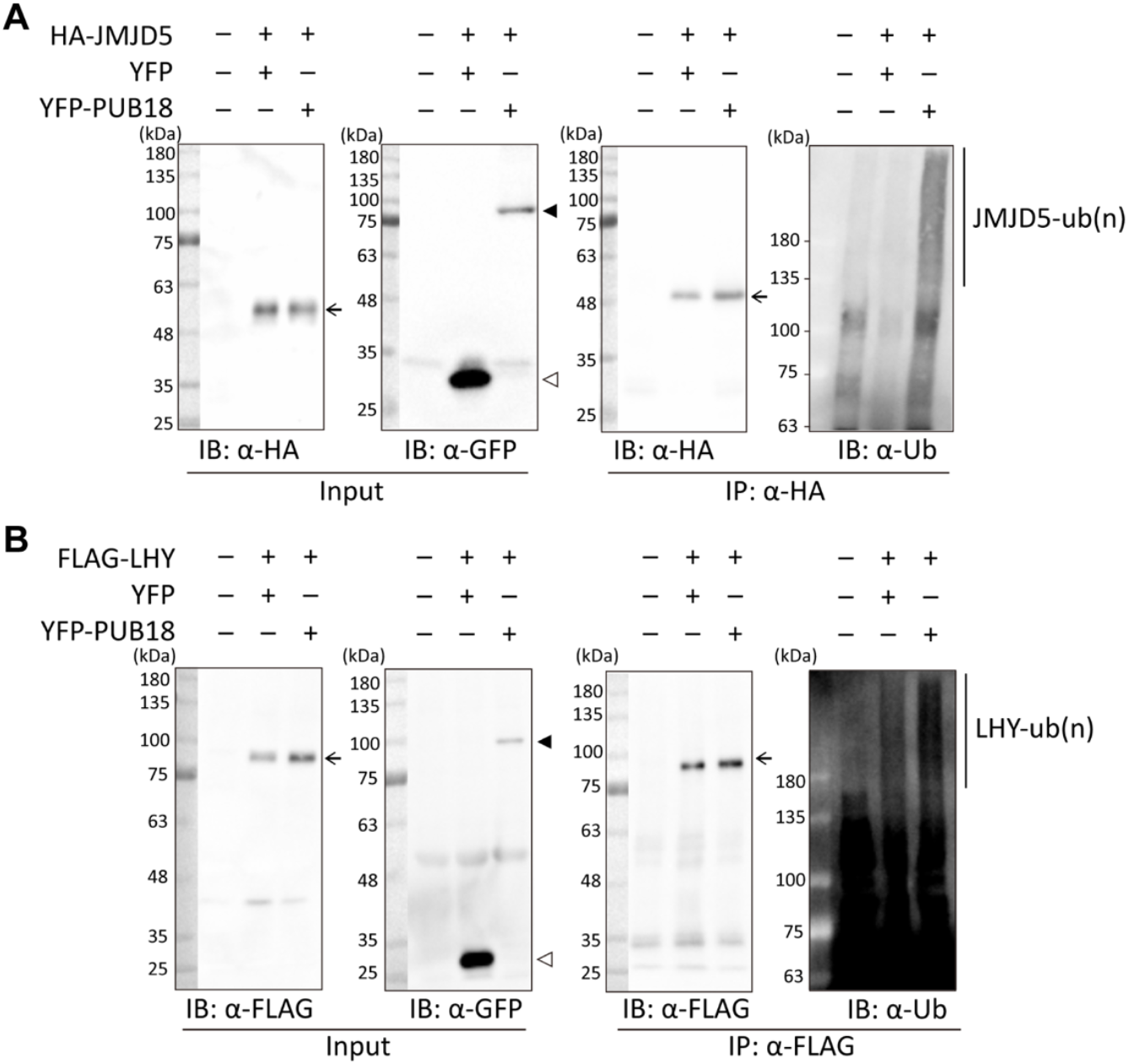
PUB18 mediates the ubiquitination of JMJD5 and LHY in plant cells. YFP-PUB18 or YFP was transiently co-expressed with **(A)** HA-JMJD5 or **(B)** FLAG-LHY in tobacco leaves to determine the ubiquitination of JMJD5 and LHY by PUB18. The HA-JMJD5 and FLAG-LHY were immunoprecipitated with anti-HA or anti-FLAG antibodies, and the ubiquitination forms were detected with anti-Ub antibodies. The arrows, black triangles, and white triangles represent the substrates (HA-JMJD5 and FLAG-LHY), YFP-PUB18, and YFP, respectively.

PUB19 is a close homolog of PUB18 in Arabidopsis (Mudgil et al. 2004), but it was not included in our E3 decoy library. We first determined whether PUB18 and PUB19 can interact with each other. In the yeast-two hybrid assays, the decoys of PUB18 and PUB19 showed interactions (**Supplemental Figure 1**). Furthermore, we conducted BiFC assay to examine whether PUB19 can also interact with LHY or JMJD5 (**Supplemental Figure 2**). However, no interaction was detected with the full-length (FL) or decoy of PUB19 (PUB19dU). In contrast, the PUB19dU showed interactions with a previously validated interactor, INDUCER OF CBF EXPRESSION 1 (ICE1) (**Supplemental Figure 2;** Wang et al., 2025), which ruled out the possibility that the dU forms of PUB19 are not expressed in tobacco leaves. Taken together, LHY, JMJD5, and CK2 interact with PUB18 but not PUB19 in the tobacco leaves.

### PUB18 and PUB19 are functionally redundant in regulating the circadian clock

LHY and JMJD5 play important roles in regulating the central circadian clock (Schaffer et al., 1998; Lu et al., 2011). As we identified PUB18 interacting with and ubiquitinating these two clock regulators, we determined whether the circadian rhythms were altered in the *pub18* mutant. Intriguingly, the circadian expression of *CCA1* in the *pub18* (25.96 ± 0.74 hr) was similar to the wildtype (25.77 ± 0.11 hr) (**Figure 6**). As *PUB18* and *PUB19* have been reported to exhibit redundant functions in abiotic stress responses (Seo et al., 2012), we generated *pub18/pub19* double mutants to examine their circadian clock period together with their single mutants. The results showed that the *pub18/pub19* double mutant displayed a significantly shorter period (25.01 ± 0.97 hr) compared to the wild type (26.77 ± 0.11 hr), while the clock rhythms of *pub19* (25.96 ± 0.29 hr) was not distinguishable from wildtype. These findings indicated that PUB18 and PUB19 act redundantly in regulating the circadian clock.

**Figure 6.**
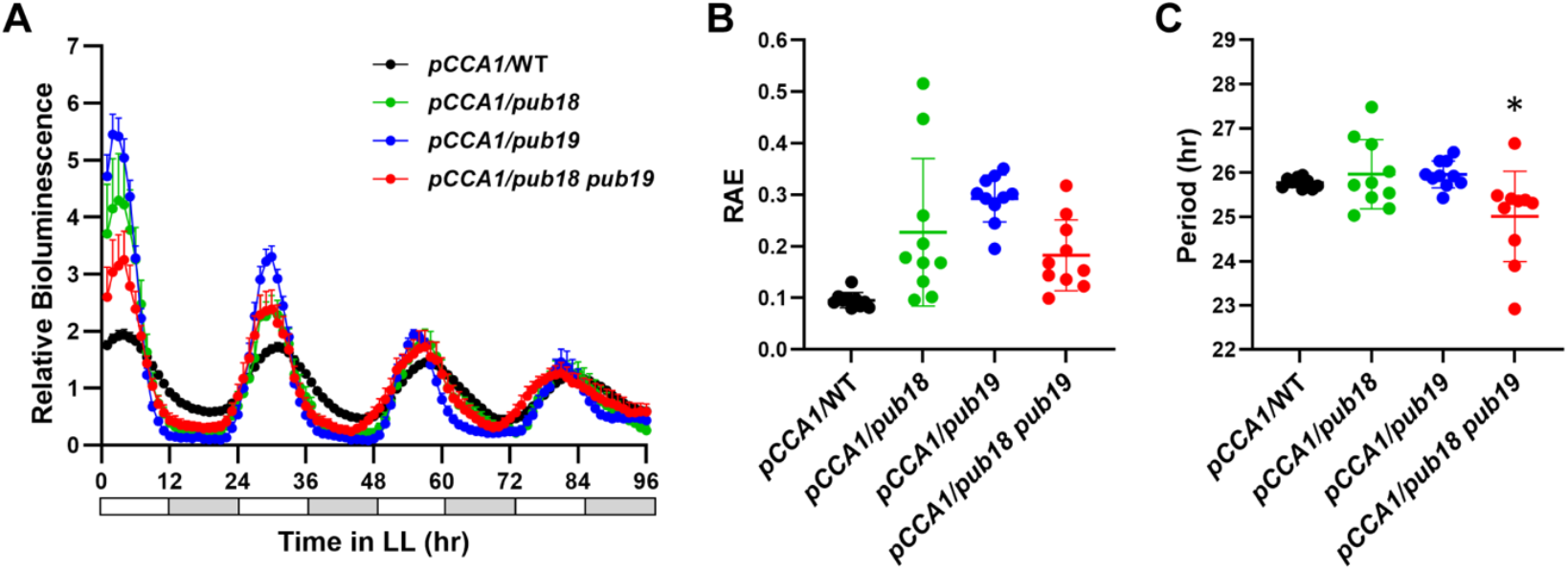
PUB18 and PUB19 are redundant to maintain circadian rhythm. The bioluminescence assays to measure the *pCCA1::LUC2* rhythms in WT, *pub18, pub19*, and *pub18*/*pub19* seedlings. The seedlings were entrained under 12 hr light /12 hr dark for 8 days before being transferred to constant light for the detection of **(A)** autoluminescence signal every hour. The **(B)** relative amplitude error (RAE) and **(C)** period length were analyzed using Fast Fourier Transform-Nonlinear Least Squares (FFT-NLLS). Error bars indicate SEM (n=10 for each line), and an asterisk indicates the statistical significance compared with the WT (t-test; \**p* < 0.05).

## Discussion

Protein ubiquitination plays a crucial role in regulating multifaceted biological pathways. Recent proteomic studies have identified 6453 ubiquitinated proteins in Arabidopsis; however, the confirmed E3-substrate pairs remain limited (Song et al., 2024). Here, we established a streamlined platform and cost-effective screen protocol to systematically identify the E3-substrate pairs and showcased this with the identification of potential E3s for the Arabidopsis clock regulators (**Figures 1 and 2**). Further, we demonstrated that the identified PUB18 interacts and ubiquitinates JMJD5 and LHY clock regulators and affects circadian clock rhythmicity (**Figure 3B, 4, 5, and 6**).

The identification of E3–substrate complexes is technically and genetically challenging due to the rapid degradation of ubiquitinated substrates and the high genetic redundancy of E3s. Our E3 screen platform was designed to overcome these obstacles while remaining feasible in a standard laboratory setting. First, because E3–substrate interactions are often unstable and may lead to false negatives in traditional yeast two-hybrid assays, we constructed a library using decoys rather than full-length E3s. Second, we adopted a high-throughput format by maintaining yeast cells that each express a single E3 decoy in individual wells of a 96-well plate. This design ensures that every E3– substrate pair can be tested without growth competition between strains, which could otherwise obscure the identification of interactors, particularly for slow-growing E3-expressing yeast. The one-to-one interaction format also eliminates the need for labor-intensive sequencing typically required in pooled library screens. Moreover, yeast mating and spotting can be easily performed using multichannel pipettes and 96-well plates, avoiding the requirement for robotics. The entire screening process, from generating diploid yeast cells through the first round of testing, can be completed within 1 to 1.5 weeks, and multiple substrates can be assayed in parallel. While this approach effectively narrows down E3 candidates, we acknowledge that the initial rapid screen may yield false positives. To address this, we included a secondary confirmation step using serial dilution assays, which requires only an additional week to complete. Together, these features establish a practical and scalable platform for systematically uncovering E3–substrate interactions, providing a valuable resource for dissecting ubiquitination-mediated regulatory networks.

We selected 21 clock regulators for the pilot screen and identified 56 E3 decoys that potentially interact with 16 of them. Nine of these regulators, including TOC1, CCA1, RVE4/6/8, BOA, LWD1, GI, and JMJD5, have previously been reported as ubiquitinated targets (Song et al., 2024; Lee and Feke et al., 2018), yet only a few E3s have been shown to alter circadian rhythms (Mas et al., 2003; Kiba et al., 2007; Yu et al., 2008). Our findings, therefore, substantially expand the list of candidate E3s that may directly regulate circadian clock components (**Figure 2**). Among these candidates, the F-box E3s ACF1 and CFK1 have been implicated in clock function, although their mechanisms in Arabidopsis remain unknown (Feke et al., 2019). CFK1 is a component of the COP9 signalosome (CSN), a functional ubiquitin ligase complex required for circadian regulation in *Neurospora* and *Drosophila* (He et al., 2005; Knowles et al., 2009; Franciosini et al., 2013). Our interaction network revealed that ACF1 interacts with RVE6 and CFK1 with PRR9, suggesting that these E3s may contribute to circadian regulation through modulation of these targets.

The interaction network also suggests more complex co-regulatory relationships. For example, GI is known to be regulated by the F-box protein ZTL and the RING-type E3 COP1 (Kim et al., 2013; Yu et al., 2008), and we additionally identified PUB25 and EIN3 BINDING F-BOX2 (EBF2) as GI interactors (**Figure 2**). ZTL and COP1 regulate GI in response to light and temperature cues (Kim et al., 2007; Kim et al., 2020; Jang et al., 2015), PUB25/26 modulate the stability of cold regulators (Wang et al., 2023), and EBF1/2 and SDIR1 promote the ethylene response to temperature fluctuations (Hao et al., 2021). GI integrates photoperiodic and temperature signals to generate thermomorphogenic rhythms, thereby promoting environmental adaptation (Park et al., 2020). These findings suggest that ZTL, COP1, EBF2, and PUB25 may collectively fine-tune GI to integrate multiple environmental signals. Moreover, our finding that ZTL interacts with LWD1, a transcriptional activator that promotes CCA1 expression at dawn (Wu et al., 2016), indicates that ZTL may regulate the clock not only through GI but also through LWD1. Our systematic identification of E3–substrate interactions using the E3 decoy library platform expands the Arabidopsis circadian interactome and provides new hypotheses for how ubiquitination fine-tunes the integration of environmental signals into the clock. This platform can be a valuable community resource to dissect the E3 functions and elucidate the mechanisms of protein ubiquitination in various physiological processes.

## Materials and Methods

### Construction of the Arabidopsis E3 Decoy Ubiquitin Ligase Yeast Screen Platform

Arabidopsis E3 decoys from the GATEWAY *pENTR* library (Feke et al., 2019) were subcloned into yeast two-hybrid *pDEST32* vectors (Thermo Scientific, cat#PQ1000101), fusing them with an N-terminal GAL4-DNA binding domain (BD) using Gateway recombination-based cloning (Thermo Scientific, cat#11791020). The constructs were verified by sequencing and then transformed into the yeast AH109 strain using the Frozen-EZ Yeast Transformation II™ kit (ZYMO RESEARCH, cat#T2001). Yeast colonies expressing the pDEST32-E3 decoy were selected on minimal medium plates (BioShop, cat#YNB406) containing 2% glucose and Leucine dropout amino acid supplement (SD-Leu). For each pDEST32-E3 decoy yeast clone, a single colony was cultured in liquid SD-Leu medium at 30°C and stored in 25% glycerol at -80°C for long-term storage. The E3 decoy yeast library consists of 283 E3 decoys (**Supplemental Table 1**), with each clone placed in a separate well across three 96-well plates, creating a high-throughput platform for protein interaction analysis.

### Cloning of Clock Regulators as Baits for Yeast Screen

Some clock regulators, fused with the N-terminal GAL4-active domain (AD), were obtained from the Arabidopsis transcription factor ORF collections (Pruneda-Paz et al., 2014) and purchased from ABRC (https://abrc.osu.edu/). These included *CIRCADIAN CLOCK ASSOCIATED 1* (*CCA1*, DEST-U21-C11), *LATE ELONGATED HYPOCOTYL* (*LHY*, DEST-U01-G05), *PSEUDO-RESPONSE REGULATOR 3* (*PRR3*, DEST-U11-D05), *PRR9* (DEST-U18-H03), *PRR7* (DEST-U21-C10), *TIMING OF CAB EXPRESSION 1* (*TOC1*, DEST-U21-C12), *REVEILLE6* (*RVE6*, DEST-U13-F10), *RVE8* (G19901), *BROTHER OF LUX ARRHYTHMO* (*BOA/NOX*, DEST-U05-G10), *JUMONJI DOMAIN CONTAINING 5* (*JMJD5*, DEST-U09-G11), *FLOWERING BASIC HELIX-LOOP-HELIX 1* (*FBH1*, DEST-U21-D01), *TEOSINTE BRANCHED 1-CYCLOIDEA-PCF20* (*TCP20*, DEST-U15-G05), *CCA1 HIKING EXPEDITION* (*CHE*, G13511), *NIGHT LIGHT-INDUCIBLE AND CLOCK-REGULATED 1* (*LNK1*, U88255), *CK2* (GC105364), and *HEAT SHOCK FACTORS B2b* (*HSFB2b*, DEST-U04-G04). For additional clock regulators, such as *RVE4, GIGANTEA* (*GI*), *LIGHT-REGULATED WD1* (*LWD1*), *LWD2*, and *TCP22*, their coding sequences were amplified from cDNA, cloned into the *pENTR-TOPO-D* vector (Thermo Scientific, cat#K240020SP), and then subcloned into the *pDEST22* vector (Thermo Scientific, cat#PQ1000101) with an N-terminal GAL4-active domain (AD) fusion. All *pDEST22* yeast expression vectors were transformed into yeast strain Y187, and single colonies were selected on SD-Trp plates.

### High-Throughput Yeast Two-Hybrid Screens

To examine interactions between the clock regulators and the E3 decoy library, diploid hybrids of *pDEST22-clock regulators* or *pDEST22* empty vector control in Y187 and *pDEST32-E3 decoys* in AH109 were generated by yeast mating. Briefly, the *pDEST22* vector culture in Y187 was mixed with the *pDEST32-E3 decoy* culture in AH109, supplemented with YPDA medium (10 g yeast extract, 20 g peptone, 20 g glucose, and 0.2 g adenine per liter) in 96-well plates. After overnight incubation (16 to 24 hours) at 30 °C, each diploid culture in the 96-well plates was plated on SD-Leu-Trp and SD-Leu-Trp-His medium plates to select for potential interactions. The growth of diploids containing the *pDEST22* empty vector and *pDEST32-E3 decoys* on the selective plates served as background controls for determining protein-protein interactions.

### Verification of Protein Interactions with Yeast Two-Hybrid Assays

To verify the interactions between E3 decoy candidates and clock regulators, a yeast two-hybrid assay with serial dilution was performed. Cultures of *pDEST22-CCA1, pDEST22-LHY, pDEST22-JMJD5*, and *pDEST22-CK2* were mated with the respective potential *pDEST32-E3 decoy* candidates. The resulting diploid yeast cells were plated on SD-Leu-Trp medium. After 2 days of incubation at 30 °C, a single colony was selected and cultured in SD-Leu-Trp liquid medium. The culture was incubated overnight (16 to 24 hours) at 30 °C, and the optical density (OD) was adjusted to 0.4 (1x). The cultures were then further diluted to 10x and 100x concentrations. A 2.5 μL aliquot of each culture was plated onto SD-Leu-Trp and SD-Leu-Trp-His medium plates, followed by incubation at 30 °C for 3 to 10 days.

### Bimolecular Fluorescence Complementation (BiFC) assay

The coding regions of *LHY, JMJD5*, and *CK2* were subcloned into *pEarleyGate201-YN*, while *PUB18* and *PUB19* were subcloned into *pEarleyGate202-YC* (Lu et al., 2010) to generate YFP N-terminus or C-terminus fusion constructs. The resulting constructs were then transformed into the Agrobacterium GV3101 strain. The *pEarleyGate202-NLS-mCherry* construct was co-expressed as a nuclear marker, and empty vectors were used as negative controls. The overnight Agrobacterium culture was pelleted, resuspended in infiltration buffer (5% sucrose, 0.1% glucose, 0.1% MgSO4, 0.1‰ Silwet, and 0.45 mM acetosyringone), and the optical density (OD) was adjusted to 0.5 at 600 nm. The mixture was kept on ice for 1 hour before infiltration into tobacco leaves. After 2 days, fluorescence signals were observed using an Apotome.2 (Zeiss).

### Co-immunoprecipitation (co-IP) assay

The coding sequences of *PUB18* and *PUB19* were amplified from genomic DNA of Col-0 seedlings and cloned into the *pCRTM8/GW/TOPO* entry vector (Thermo Scientific, cat#K250020). The LR Clonase II enzyme mix (Thermo Scientific, cat#11791020) was used to transfer the inserts into the destination vectors *pEarleyGate 202* (Earley et al., 2006) and *pUBQ10::YFP-GW* (Michniewicz et al., 2015). The *pEarleyGate 202*-*LHY, pEarleyGate 202*-*JMJD5* or *pUBQ10::YFP-PUB18* constructs were transformed into Agrobacterium GV3101 and transiently co-expressed in tobacco leaves using Agrobacterium infiltration. After 2 days, tobacco leaves were harvested, frozen in liquid nitrogen, and ground. Proteins were extracted using Tris-protein extraction buffer (50 mM Tris-HCl, pH 7.4, 150 mM NaCl, 10% glycerol, 1% NP-40, 1 mM PMSF, 1× protease inhibitor cocktail (Roche, cat#11697498001), 5 μM EDTA, pH 8.0). The protein extracts were incubated at 4°C for 3 hours with GFP-Trap beads (Chromo Tek, cat#gtma). The immunoprecipitated samples were heated at 95°C for 5 minutes in SDS sample buffer (25 mM Tris-HCl, pH 6.8, 2% SDS, 5% glycerol, 12.5 mM EDTA, 0.5% β-mercaptoethanol, 0.1% (w/v) bromophenol blue). Protein separation was performed by SDS-PAGE, followed by immunodetection using anti-FLAG (Sigma, cat#F1804) and anti-GFP (Abcam, cat#ab-290) antibodies. The signal was detected using the iBright™ imaging system (Invitrogen).

### Measurement of Circadian Rhythms

Arabidopsis plants carrying the *pCCA1::LUC2* reporter in the Col-0 background (Wu et al., 2008) were crossed with *pub18* (*SALK_001831*), *pub19* (*SALK_058791*), or *pub18/pub19* mutants (Seo et al., 2012) obtained from ABRC. The seedlings were grown on 1/2 MS medium without sucrose under a 12-hour light/12-hour dark cycle at 22°C for 8 days before being transferred to a 96-well black plate. Prior to imaging, 1/2 MS liquid medium containing 0.5 mM D-Luciferin (GoldBio, cat# LUCK-100) was added to each well. The plate was then incubated under constant light at 22°C in a Phenotron Pro system (Taiwan Hipoint) to record the autoluminescence signals every hour. The luminescence signal of each seedling was quantified as the mean intensity and the data were imported into the Biodare2 platform (biodare2.ed.ac.uk; Zieliński et al., 2014; Zieliński et al., 2022) for analysis of period length and amplitude using Fast Fourier Transform-Nonlinear Least Squares (FFT-NLLS).

### Accession numbers

Sequence data from this article can be found in The Arabidopsis Information Resource under the following accession numbers: *PUB18* (AT1G10560), *PUB19* (AT1G60190), *LHY* (AT1G01060), *JMJD5* (AT3G20810), *CK2* (AT3G50000), *CCA1* (AT2G26830), *RVE8* (AT3G09600), *RVE4* (AT5G02840), *RVE6* (AT5G52660), *PRR3* (AT5G60100), *PRR7* (AT5G02810), *PRR9* (AT2G46790), *BOA* (AT5G59570), *LWD1* (AT1G12910), *LWD2* (AT3G26640), *TCP20* (AT3G27010), *TCP22* (AT1G72010), *TOC1* (AT5G61380), *CHE* (AT5G08330), *LNK1* (AT5G64170), *FBH1* (AT1G35460), *GI* (AT1G22770), *HSFB2b* (AT4G11660).

## List of supplemental data

**Supplementary Table 1.** The list of Arabidopsis E3 decoy yeast library for the high-throughput yeast two-hybrid screens.

**Supplementary Table 2.** The List of clock regulators for high-throughput E3 Decoy yeast two- hybrid screens.

**Supplementary Table 3.** The E3 decoy-clock regulator interaction pairs from the high-throughput yeast two-hybrid screen.

**Supplementary Table 4.** The primers were used in this study.

## Acknowledgments

We thank Dr. Joshua Gendron for generously providing the E3 decoy collection and Dr. Shu-Hsing Wu for the *pCCA1::LUC+* transgenic line. We are also grateful to Dr. Shu-Hsing Wu and Dr. Huang-Lung Tsai for their valuable discussions. We appreciate the excellent technical assistance and support from the Technology Commons, College of Life Science, National Taiwan University, which greatly facilitated our research.

## Author Contributions

The study was conceptualized by C.M.L. and J.M.G., and the manuscript was written by Y.T.T., C.A.C. and C.M.L.. Y.Y.T., C.A.C. and C.M.L. designed, executed the experiments and analyzed the data. J.M.G. provided the E3 decoy collection in the *pENTR* vectors.

## Data availability

The Arabidopsis E3 decoy library will be deposited at the Arabidopsis Biological Resource Center (ABRC) as a community resource. The step-by-step protocol will be available on https://www.protocols.io/.

## Funding

C.M.L.: MOST-109-2311-B-002-030-MY2, NSTC-111-2311-B-002-012-MY3, NTU-CC-112L891804, and NTU-CC-114L895003

Y.T.T.: MOST-110-2811-B-002-523-, NTU-CC-111L4000, NTU-CC-112L4000, NTU-CC-113L4000, and NSTC-113-2811-B-002-122-

C.A.C.: NSTC PhD fellowship and NTU Outstanding Doctoral Students Scholarship

---

**Supplemental Figure 1.**
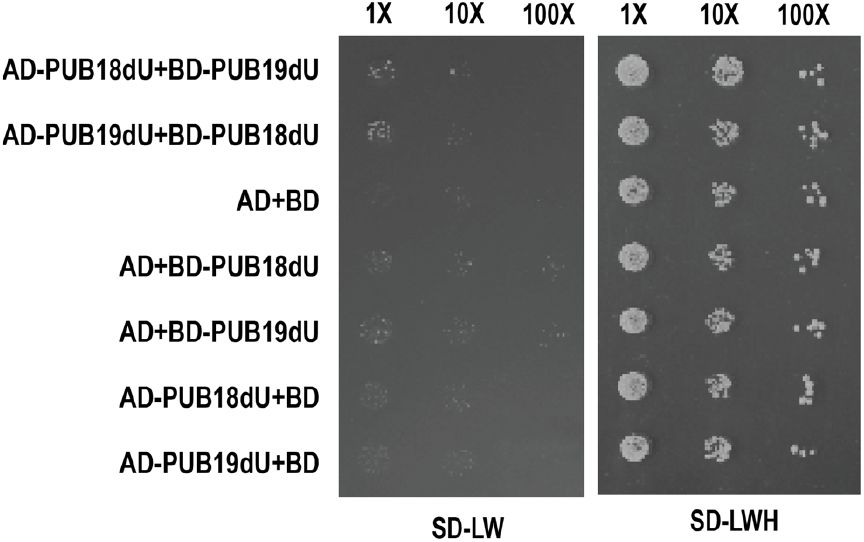
The yeast two-hybrid assays were used to examine the interactions between the decoy form of PUB18 and PUB19. The U-box deletion (dU) forms of PUB18 and PUB19 were fused with GAL4-AD in the *pDEST22* vector or GLA4-BD in the *pDEST32* vectors to test the interactions with the yeast two-hybrid assays. The 10-fold (10X) and 100-fold (100X) dilutions were made from yeast culture with O.D.= 0.4 (labeled with 1X). The yeast culture was grown on SD-Leu-Trp (SD-LW) or SD-Leu-Trp-His (SD-LWH) medium for interaction tests.

**Supplemental Figure 2.**
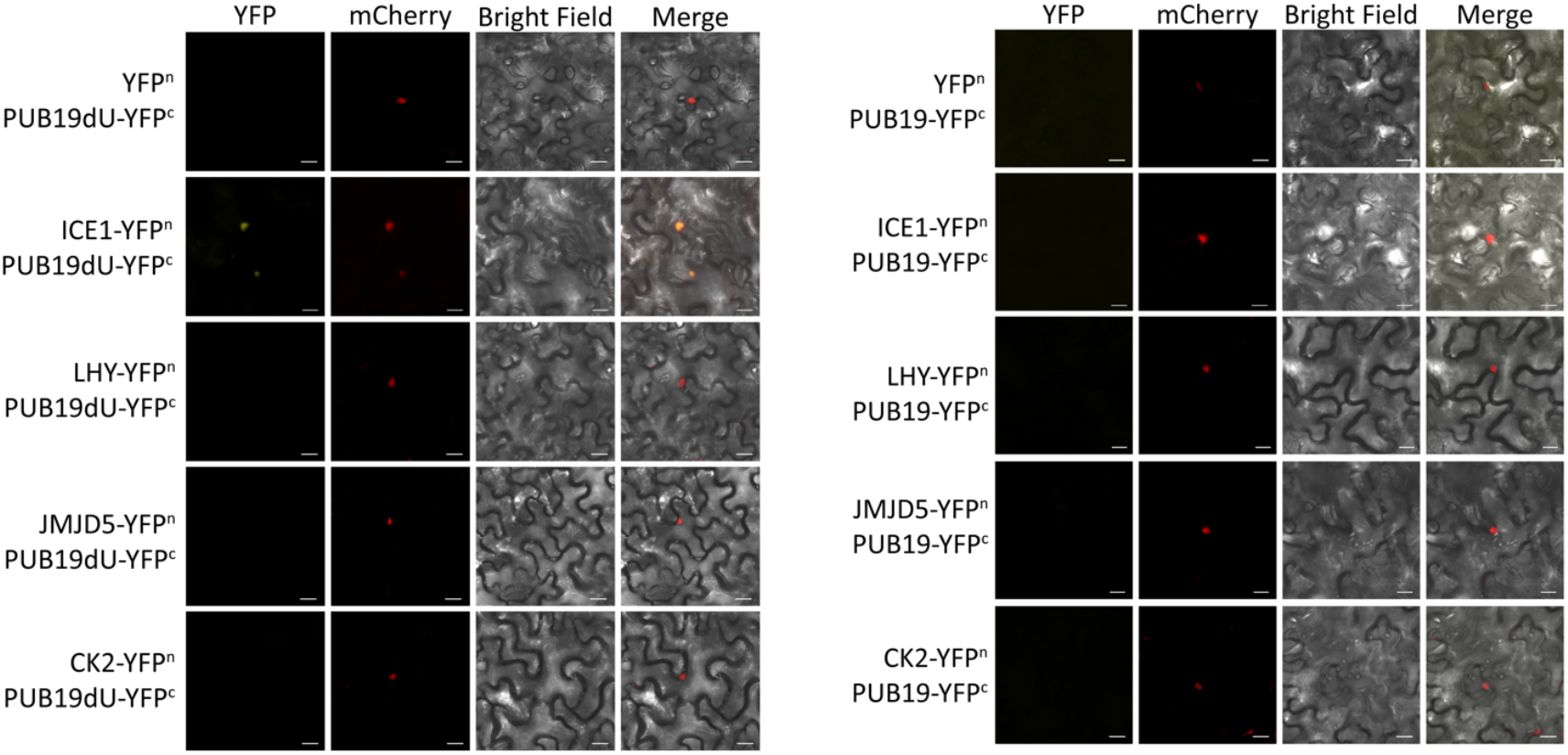
JMJD5 and LHY didn’t interact with PUB19 and PUB19dU in *N. benthamiana*. The Bimolecular fluorescence complementation (BiFC) assays to examine the interactions between JMJD5 and LHY fused to the N-terminal of YFP (YFP^n^) and full-length PUB19 (PUB19) or U-box-deleted PUB19 (PUB19dU) fused to the C-terminal of YFP (YFP^c^) in tobacco leaves. The NLS-mCherry was co-expressed as a nuclear marker.Scale bar= 20µm.

